# Various evolutionary trajectories lead to loss of the tobramycin-potentiating activity of the quorum sensing inhibitor baicalin hydrate in *Burkholderia cenocepacia* biofilms

**DOI:** 10.1101/409078

**Authors:** Andrea Sass, Lisa Slachmuylders, Heleen Van Acker, Ian Vandenbussche, Lisa Ostyn, Aurélie Crabbé, Laurent Chiarelli, Silvia Buroni, Filip Van Nieuwerburgh, Emmanuel Abatih, Tom Coenye

## Abstract

Combining antibiotics with potentiators that increase their activity is a promising strategy to tackle infections caused by antibiotic-resistant and -tolerant bacteria. As these potentiators typically do not interfere with essential processes of bacteria, it has been hypothesized that they are less likely to induce resistance than conventional antibiotics. However, evidence supporting this hypothesis is lacking. In the present study, we investigated whether *Burkholderia cenocepacia* J2315 biofilms develop resistance towards one such adjuvant, baicalin hydrate (BH), a quorum sensing inhibitor known to increase antibiotic-induced oxidative stress. Biofilms were repeatedly and intermittently treated with tobramycin (TOB) alone or in combination with BH for 24 h. After each cycle of treatment, the remaining cells were quantified using plate counting. After 15 cycles, biofilm cells were less susceptible to treatments with TOB and TOB+BH, compared to the start population, and the potentiating effect of BH towards TOB was lost. Whole genome sequencing was performed to probe which changes were involved in the reduced effect of BH and mutations in 14 protein-coding genes were identified (including mutations in genes involved in central metabolism and in BCAL0296, encoding an ABC transporter), as well as a partial deletion of two larger regions. No changes in the minimal inhibitory or minimal bactericidal concentration of TOB or changes in the number of persister cells were observed in the evolved populations. However, basal intracellular levels of reactive oxygen species (ROS) and ROS levels found after treatment with TOB were markedly decreased in the evolved populations. In addition, in evolved cultures with mutations in BCAL0296, a significantly reduced uptake of TOB was observed. Our results indicate that resistance towards antibiotic-potentiating activity can develop rapidly in *B. cenocepacia* J2315 biofilms and point to changes in central metabolism, reduced ROS production, and reduced TOB uptake as potential mechanisms.

**Importance:** Bacteria show a markedly reduced susceptibility to antibiotics when growing in a biofilm, which hampers effective treatment of biofilm-related infections. The use of potentiators that increase the activity of antibiotics against biofilms has been proposed as a solution to this problem, but it is unclear whether resistance to these potentiators could develop. Using an experimental evolution approach, we convincingly demonstrate that *Burkholderia cenocepacia* biofilms rapidly develop resistance towards the tobramycin-potentiating activity of baicalin hydrate. Whole genome sequencing revealed that there are different mechanisms that lead to this resistance, including mutations resulting in metabolic changes, changes in production of intracellular levels of reactive oxygen species, and differences in transporter-mediated tobramycin uptake. Our study suggests that this form of combination therapy is not ‘evolution-proof’ and highlights the usefulness of experimental evolution to identify mechanisms of resistance and tolerance in biofilm-grown bacteria.

## Introduction

Due to increasing levels of antimicrobial resistance, novel strategies to tackle bacterial infections are needed and an interesting approach is the use of antibiotic adjuvants or potentiators. Potentiators are compounds with little or no antibacterial activity that interfere with bacterial resistance mechanisms and/or increase antimicrobial activity when co-administered with an antibiotic (1-5). A well-known class of antibiotic adjuvants are quorum sensing (QS) inhibitors (QSIs) (6). QSIs target the cell-density based bacterial communication network that regulates the expression of multiple virulence factors (7, 8). Whether resistance would develop towards these adjuvants is currently unknown and QSIs have long been accepted as ‘evolution-proof’: as QSIs do not target pathways essential for growth, it has been hypothesized that development of resistance would not occur (or at least would occur less frequently), due to the lack of selective pressure favouring the rise of resistant mutants (9-13). However, natural selection occurs when heritable variation provides a fitness advantage and QS disruption can affect bacterial fitness in conditions in which a functional QS system is essential (8). This was for example shown by cultivating *Pseudomonas aeruginosa* in medium with adenosine as a sole carbon source (14). As growth on adenosine depends on the production of a nucleoside hydrolase, which is positively regulated by the key QS signal receptor LasR, a functional QS system is required for the growth of *P. aeruginosa* in these conditions (15). After addition of the brominated furanone C-30 (a known QSI), growth of *P. aeruginosa* on adenosine was impaired, resulting in selective pressure and the occurrence of resistant mutants. Adding this QSI caused mutations in repressor genes of the multidrug resistance efflux pump MexAB-OrpM, which resulted in an increased resistance towards C-30 (14). In clinical isolates of cystic fibrosis (CF) patients never exposed to C-30, mutations in the same genes were found, leading to reduced susceptibility to C-30 (14, 16). Based on these results, Maeda et al speculated that any strong selective pressure can induce resistance to antivirulence compounds (14, 17). In clinical practice, these adjuvants would be co-administered with an antibiotic. This means selective pressure imposed by this antibiotic needs to be included in the experimental set up when investigating possible development of resistance towards the adjuvants (1, 2). In addition, while most evolutionary studies on the development of resistance are carried out with planktonic cells (18-20), 65-80% of all infections are thought to be biofilm-related, and biofilm-associated bacteria typically show a reduced susceptibility towards antimicrobial agents (21).

*Burkholderia cenocepacia* is an opportunistic pathogen that causes severe lung infections in people with CF, which can further develop into a life-threatening systemic infection known as the cepacia syndrome (22). Antimicrobial therapy in CF often fails due to high innate resistance of *B. cenocepacia* towards many antibacterial agents and high tolerance associated with its biofilm-lifestyle (22, 23). Previously, several adjuvants were identified that increased the activity of tobramycin (TOB) (an aminoglycoside antibiotic frequently used in CF lung infections) (24) towards *B. cenocepacia* biofilms, including the QSI baicalin hydrate (BH) (6, 25).

The goal of the present study is to evaluate whether (and how) *B. cenocepacia* J2315 biofilm cells can develop resistance towards the TOB-potentiating activity of BH. To this end we used a slightly modified form of a previously-described bead-based biofilm assay (26) in which *B. cenocepacia* J2315 cells were repeatedly and intermittently exposed to TOB, TOB+BH or a control treatment (Fig. 1).

**Fig 1.**
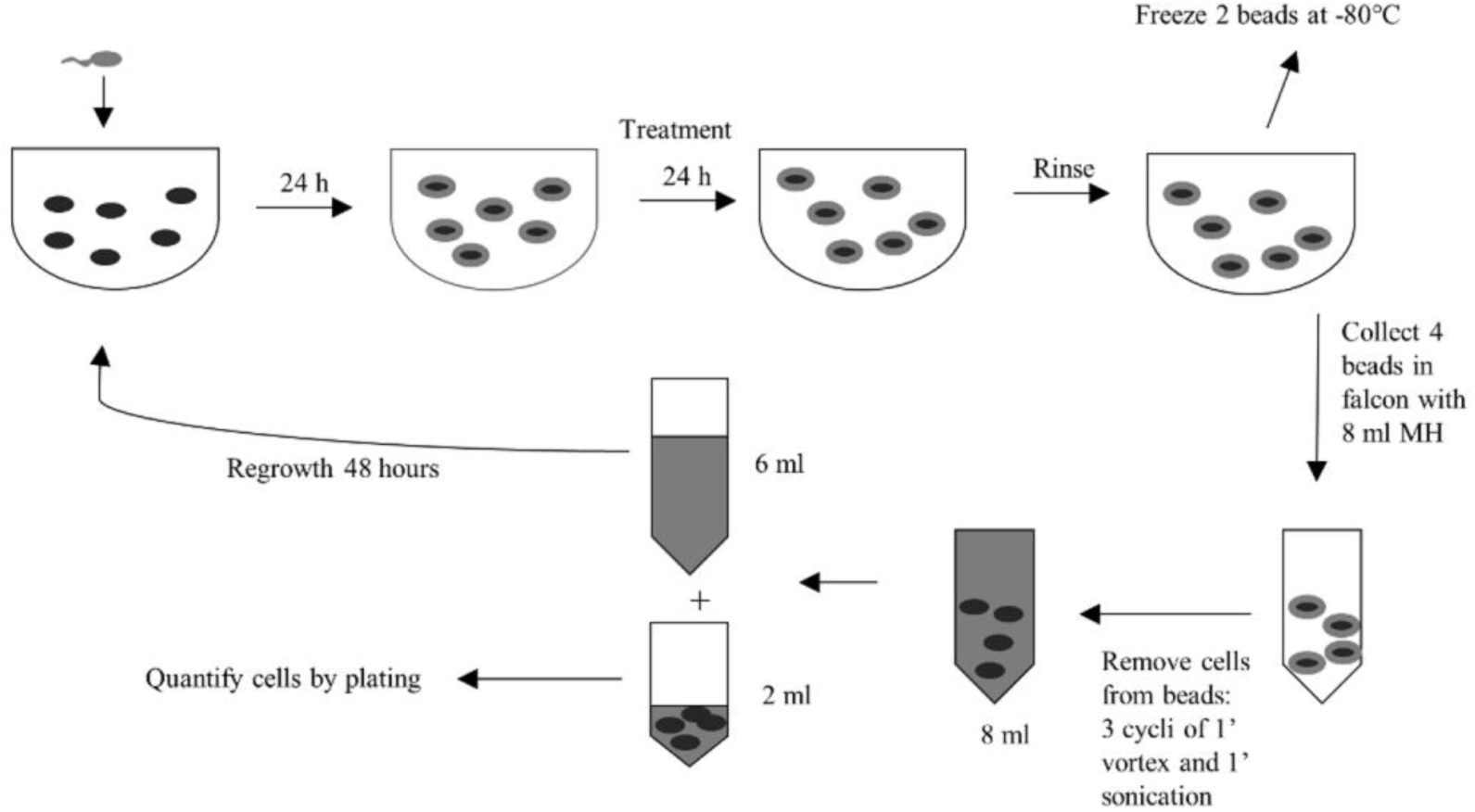
Experimental set up. Fresh inoculum (grey) is added to six cryobeads (black full circles) in each well of a 24-well microtiter plate. After 24 hours, mature biofilms (grey circles) are formed on the surface of the beads. These biofilms are treated for 24 hours. Afterwards, the supernatant is removed, and the beads are rinsed with PS. Two beads, containing a mature biofilm, are stored at - 80°C. The four other beads are transferred to a falcon tube, in which the sessile cells from the beads are harvested. A part of these cells is used for quantification, while another part is used for planktonic regrowth of the cells (48 hours).

## Results and discussion

### Experimental evolution

Three lineages of *B. cenocepacia* J2315 cells were repeatedly and intermittently exposed to TOB, TOB+BH or a control treatment (Fig. 1). After 24 h of growth on the beads, LIVE/DEAD staining was performed to evaluate biofilm formation (Fig. S1). A dense biofilm was formed in the cavity of the doughnut-shaped bead (rather than on the exterior sides of the bead) with approximately 10^7^ CFU/bead (prior to treatment).

The number of log(CFU/bead) after every cycle is shown in Fig. 2 and Table S1. After fitting a linear mixed-effect model (LMEM) (using log(CFU/bead) as the dependent variable and cycle, treatment, lineage and their two- and three-way interactions as fixed effects) and plotting the residuals against the corresponding fitted values, no departures from the main assumptions of normality and constancy of error variance were found. The remaining models were fit for each lineage separately following significance of the three way interaction effect and residuals were assessed for each of the models for each lineage. An overview of the statistical results obtained can be found in Tables 1 and S2.

**TABLE 1.** Most important results of the LMEM per lineage. Treatments were pairwise compared to TOB treatment over time.

|  | **Factor** | **t value** | **Tukey adjusted p-value** |
| --- | --- | --- | --- |
| **Lineage 1** | Untreated control | -4.09 | <0.0001 |
|  | TOB+BH | 2.71 | 0.0075 |
| **Lineage 2** | Untreated control | -4.41 | <0.0001 |
|  | TOB+BH | 6.13 | <0.0001 |
| **Lineage 3** | Untreated control | -2.49 | 0.0139 |
|  | TOB+BH | 6.08 | <0.0001 |

**Fig 2.**
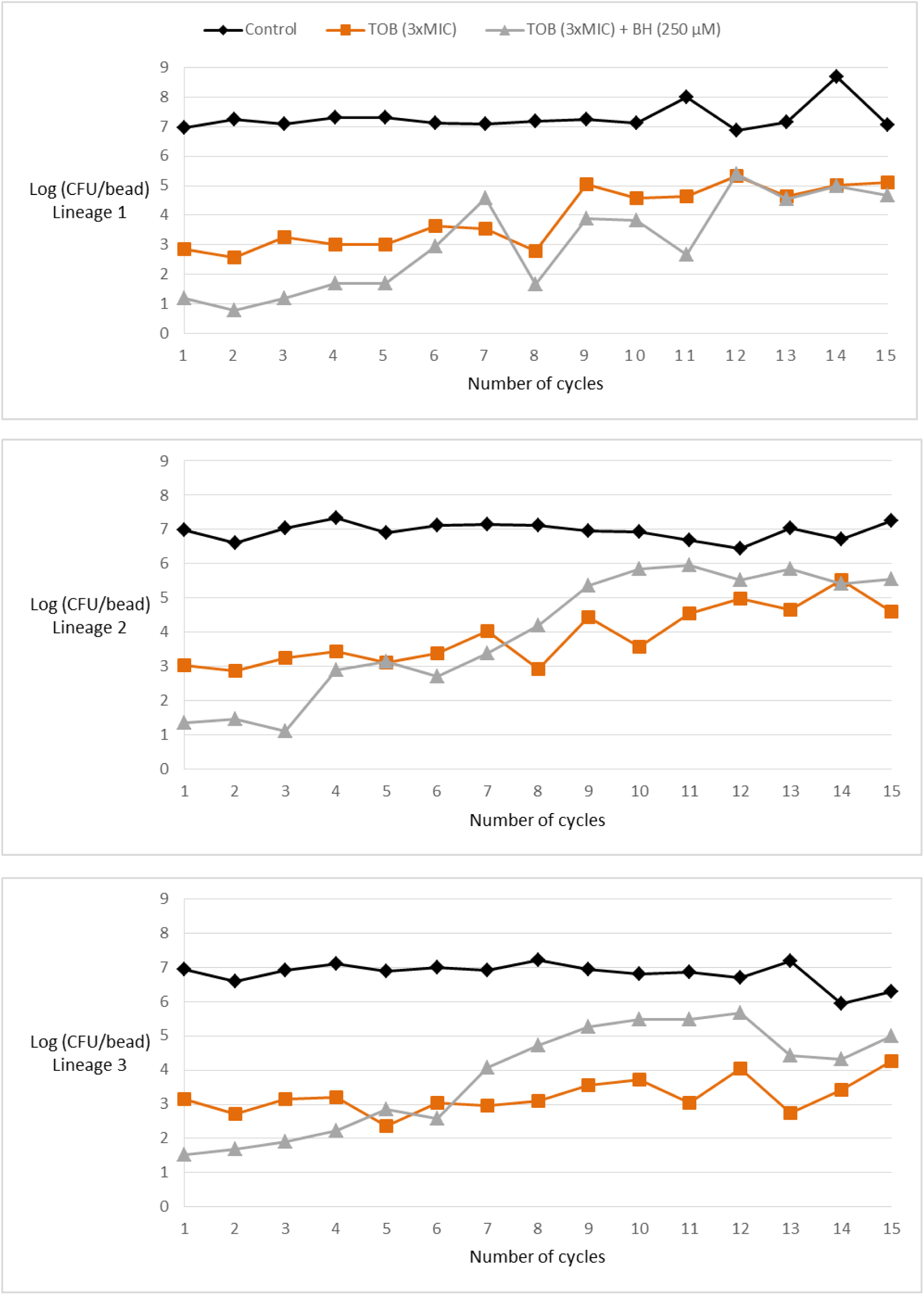
Number of *B. cenocepacia* J2315 biofilm cells, expressed as log(CFU/bead), in the untreated control (Control) and after repeated treatments with TOB or TOB+BH.

At the start of the evolution experiment, cells were more susceptible to the TOB+BH combination than to TOB alone, confirming that BH potentiates the activity of TOB against biofilms, as previously shown (6, 27). Over time, biofilm-grown *B. cenocepacia* J2315 cells became gradually less susceptible to the treatment (both to treatment with TOB alone and to the TOB+BH combination treatment); this occurred in all three lineages (Fig. 2). Evolution towards reduced susceptibility occurred significantly faster with the combined TOB+BH treatment, than for the TOB treatment (lineage 1: p = 0.0075; lineage 2 and 3: p < 0.0001) (Table 1) and our data indicate that in all lineages, the TOB-potentiating activity of BH was lost after 15 cycles, i.e. treatment with the combination TOB + BH was not able to kill more cells than treatment with TOB alone (Fig. S2).

### Genome analysis

To investigate the reason behind this decreased susceptibility, whole genome sequencing was performed. The results are summarised in Table 2. When considering all (nine) evolved lines, changes in 18 protein-coding genes were observed, as well as a partial deletion of two larger regions. Some changes were common and appeared in all evolved cultures at the same location (e.g. changes in BCAL1315, BCAL1664 and BCAM0949); we speculate these mutations were already present in the start population at low frequency and were enriched for during the experimental evolution. Other changes occurred only in one or a few samples (e.g. mutations in BCAL0929, BCAL2476a, BCAM1901) and likely arose during the evolution study. For several genes that were mutated in multiple evolved cultures we noticed the occurrence of different types of mutations (e.g. BCAL0269, BCAL1525, BCAM0965).

**TABLE 2.**
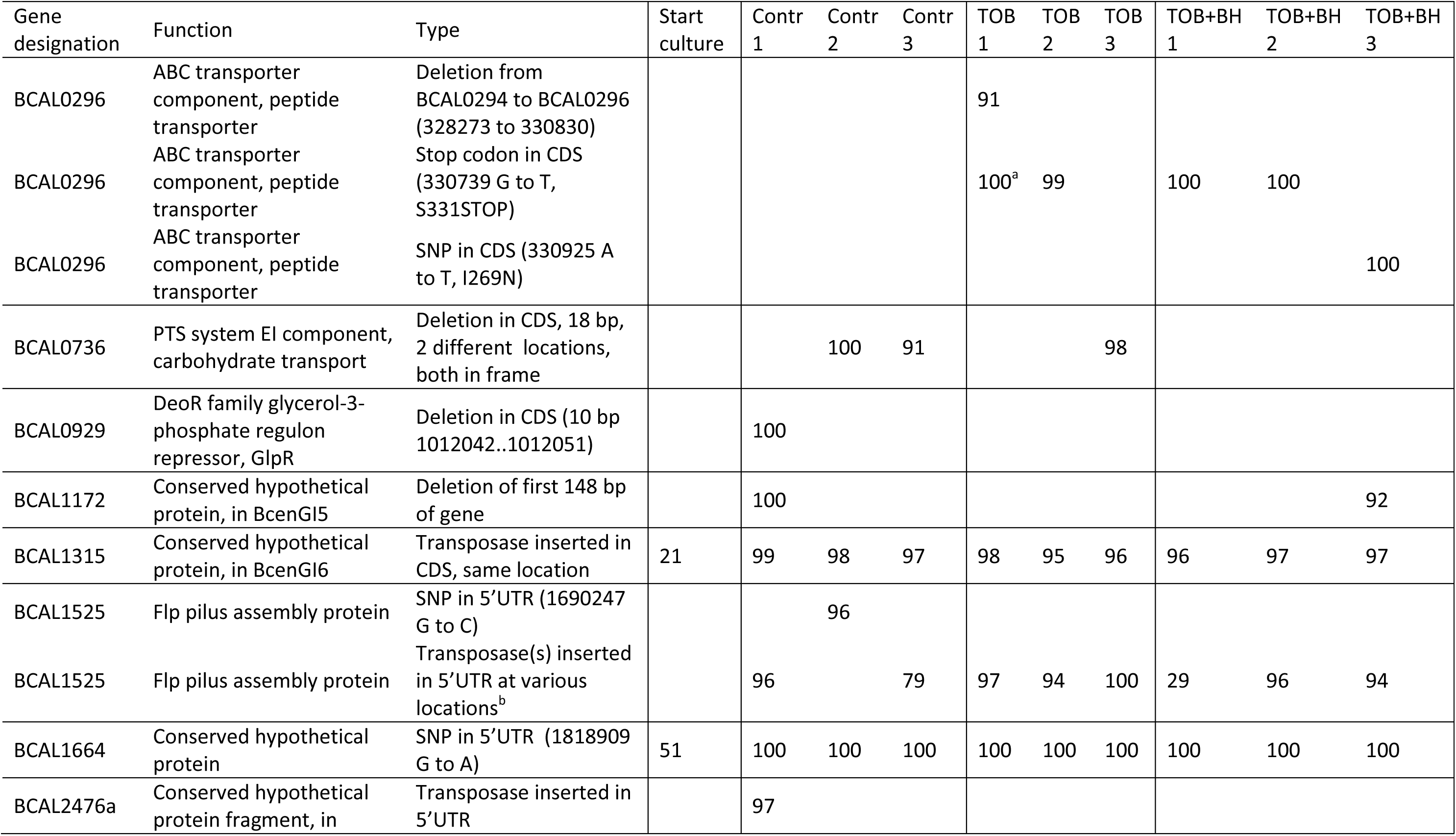
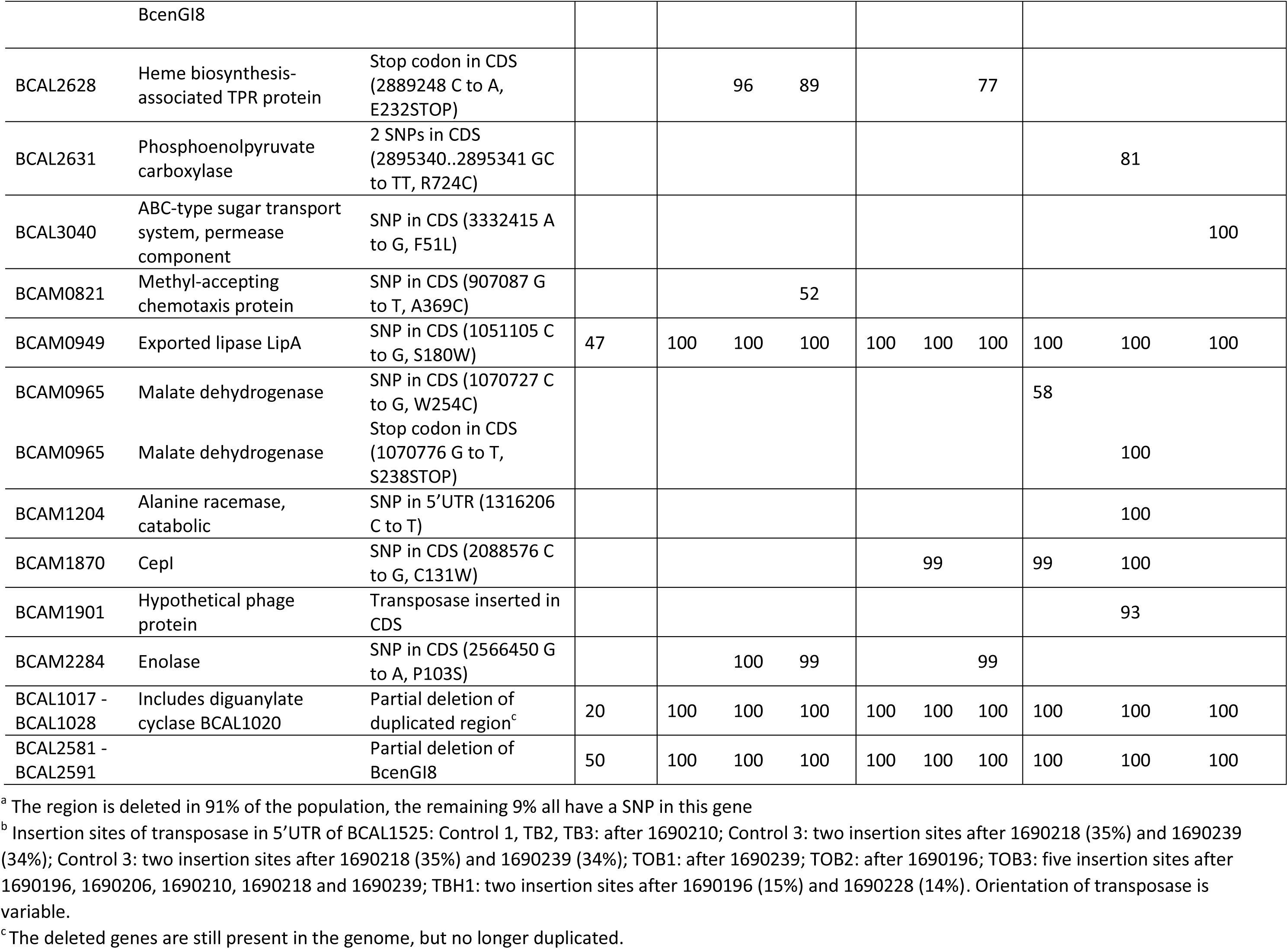
Mutations observed in evolved populations. Numbers represent variant-to-reference ratios.

All evolved cultures had mutations in the 5’ untranslated region (UTR) of BCAL1525 (either a single SNP or insertion of a transposase at variable locations, Table 2). BCAL1525 encodes an Flp pilus assembly protein and has a 307 nt long and strongly expressed 5’UTR with multiple transcription start sites (TSS) (28). BCAL1525 to BCAL1536 seem to form an operon with Flp pilus genes, although there is a terminator at position 1690709 to 1690746 between BCAL1525 and BCAL1526, and no further downstream TSS could be identified (28). Flp pili belong to the type IVb pilus family (29), which is poorly characterized in *Burkholderia* species. In *P. aeruginosa* type IV pili are thought to play a role in biofilm formation (30), although recent data suggest this may depend on the biofilm model system used (31). Mutations in BCAL1525 occur at high frequency in all evolved cultures, irrespective of the treatment, suggesting there is an evolutionary pressure to lose the pilus function in the given experimental conditions. As biofilm formation is not affected in the course of the experiment, this indicates the Flp pilus is not required for biofilm formation in these conditions.

BCAL0296 encodes for both the transmembrane and nucleotide binding domains of an ABC transport protein and it is the only gene which is mutated in all treated evolved populations (except for one evolved population treated with TOB) but not in the control evolved populations; three different types of mutations are observed in this gene (a deletion, a nonsense mutation and a non-synonymous substitution, Table 2).

Three mutated genes are related to central metabolism and occur in TOB+BH treated lineages only. BCAL2631 and BCAM0965 are both involved in oxaloacetate production. The mutation in BCAL2631 occurs only in one population exposed to TOB+BH; this gene encodes phosphoenol pyruvate kinase, which converts phosphoenol pyruvate to oxaloacetate in the ‘reverse TCA cycle’. Its activity typically results in increased oxaloacetate levels and an increased flux through the TCA cycle (32). It was designated as conditionally essential in *B. cenocepacia* J2315 in a minimal medium with only glucose as substrate (33). BCAM0965 (encoding malate dehydrogenase) is mutated in two out of three evolved populations exposed to TOB+BH. Just like phosphoenol pyruvate kinase, malate dehydrogenase (which converts malate to oxaloacetate) activity will increase cellular oxaloacetate levels, likely stimulating the TCA cycle. BCAM0965 is also conditionally essential in *B. cenocepacia* J2315 (33). On top of that, in one TOB+BH exposed evolved population, a mutation in a gene encoding the permease of a glucose/mannose ABC transporter (BCAL3040) was observed, likely affecting uptake of certain carbohydrates.

None of the mutations observed were located in genes known to be responsible for aminoglycoside resistance such as genes encoding for aminoglycoside-modifying enzymes, efflux pumps, or ribosome methyltransferases, or in genes encoding for ribosomal RNAs (34, 35). Since mutations in these genes often come at the cost of reduced relative growth fitness (36), the re-growth phase between treatment cycles might have prevented accumulation of such mutations.

### Role of QS and QSI

BH was previously described as a QSI (6), and in a recent study we showed it also has QS-independent activities, including modulating the oxidative stress response, that potentiate TOB in *B. cenocepacia* (27). To evaluate whether BH directly inhibits the *N*-octanoyl-L-homoserine lactone synthase CepI (37, 38), an enzymatic assay with purified CepI was carried out. CepI enzymatic activity was first tested at the concentration of BH slightly below that used for the evolution study (200 μM) and at this concentration BH completely inhibits CepI; subsequently, the IC_50_ was determined (46.8 ± 6.8 μM) which confirmed that BH inhibits CepI in a concentration dependent way (Fig. S3).

The mutation in BCAM1870, coding for CepI, is found in two evolved populations exposed to TOB+BH and in a single population exposed to TOB only; the same mutation (C131W) is found in these three populations. Using qPCR, we investigated the expression of *cepI* (BCAM1870) and two QS-regulated genes (*aidA* [BCAS0293] and *zmpA* [BCAS0409]) (38) in cultures derived from the TOB-exposed biofilm of lineage 2. In stationary phase expression levels of these genes were not altered, but lower expression levels for these genes were observed in late log-phase cultures of the *cepI* mutant, with remarkably low levels of *aidA* expression (approx. 60-fold lower expression compared to the control) (Fig. 3). This confirms that QS is indeed affected in this mutant. How the mutation in *cepI* contributes to an increased fitness of the evolved *B. cenocepacia* populations is currently unknown.

**Fig 3.**
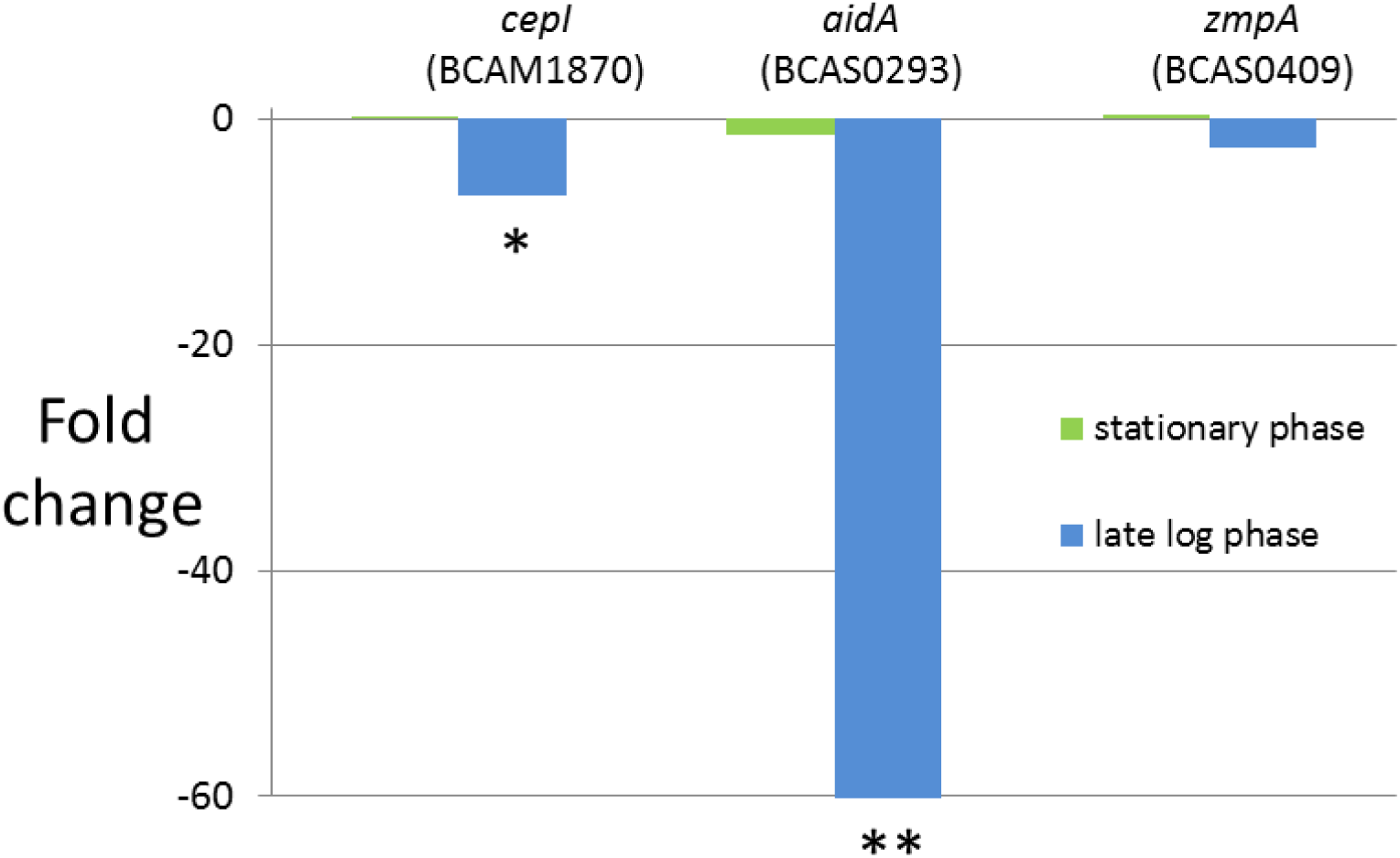
Expression of *cepI, aidA* and *zmpA* in TOB-exposed evolved lineage compared to control lineage, as determined by qPCR. Data shown are fold change in TOB-exposed lineage compared to control. ^*^, p<0.05; ^**^, p<0.01.

### Phenotypic characterisation of evolved lines

The lack of mutations in known TOB resistance genes suggested that the overall gradual decrease in susceptibility of *B. cenocepacia* J2315 biofilm cells treated with TOB alone is not caused by a resistance mechanism specific for TOB. This is in line with the MIC and MBC values for TOB obtained for the different lineages (Table 3): in the presence or absence of BH, MIC and MBC values for TOB in the evolved lines are equal to or within one 2-fold dilution of the values obtained for the start culture.

**TABLE 3.** MIC and MBC of TOB in *B. cenocepacia* J2315 recovered from different samples. MIC and MBC were determined in presence (TOB+BH) and absence (TOB) of BH.

|  | Treatment | MIC of TOB<br>(µg/ml) | MBC of TOB<br>(µg/ml) |
| --- | --- | --- | --- |
| Start culture | TOB | 256 | 256 |
|  | TOB+BH | 256 | 256 |
| Lineage 1, treated<br>with TOB | TOB | 128 | 128 |
|  | TOB+BH | 128 | 128 |
| Lineage 1, treated<br>with TOB+BH | TOB | 256 | 256 |
|  | TOB+BH | 128 | 256 |
| Lineage 2, treated<br>with TOB | TOB | 128 | 128 |
|  | TOB+BH | 128 | 128 |
| Lineage 2, treated<br>with TOB+BH | TOB | 128 | 128 |
|  | TOB+BH | 128 | 128 |
| Lineage 3, treated<br>with TOB | TOB | 256 | 256 |
|  | TOB+BH | 256 | 256 |
| Lineage 3, treated<br>with TOB+BH | TOB | 256 | 512 |
|  | TOB+BH | 512 | 512 |

Secondly, we determined whether the evolutionary changes affected the number of persister cells in treated cultures (experiments concerning persisters were carried out in two-fold with lineage 3 only). Persisters are known to occur in *B. cenocepacia* biofilms (39) and mutations leading to an increased fraction of persisters could (at least partially) explain the reduced effect of TOB+BH in the evolved lineages. However, the fraction of persisters recovered from *B. cenocepacia* J2315 biofilms after exposure to 4xMIC TOB were low and very similar for the control (0.0224%), the culture evolved in the presence of TOB (0.0393%) and the culture evolved in the presence of TOB+BH (0.0548%), ruling out increased persister formation as a source for the diminishing effect of TOB+BH (Fig. S4).

Planktonic growth rate was not affected in any of the lineages evolved in the presence of TOB, but a slightly reduced growth rate was observed for lineages 1 and 2 evolved in the presence of TOB+BH (Fig. S5). This reduction in growth was most pronounced for lineage 2, particularly in early exponential and stationary growth phase. Interestingly, this is also the only lineage that has mutations in two conditionally essential genes (BCAL2631 and BCAM0965) (33), potentially explaining this growth phenotype.

Subsequently, we investigated whether there were differences in production of reactive oxygen species (ROS) between the start culture and the evolved populations. We have previously shown that bactericidal antibiotics (including TOB) induce ROS in *B. cenocepacia* biofilms and that this contributes to the antibiotic-mediated killing in *B. cenocepacia* (39-41). In addition, we have previously shown that BH increases TOB-induced oxidative stress in a QS-independent way (27). ROS are an inevitable by-product of aerobic respiration and as we observed mutations in genes involved in oxaloacetate production (BCAL2631 and BCAM0695) or glucose/mannose transport (BCAL3040) in all lineages treated with TOB+BH (but not in lineages treated with TOB alone or in control lineages, Table 2) we hypothesised these mutations could affect ROS levels. First, we investigated if basal ROS levels (i.e. ROS levels observed in the absence of a treatment with a bactericidal antibiotic) were different between the start culture and the evolved populations. For the control populations and populations treated with TOB, significantly increased basal ROS levels were observed in two and one of the lineages, respectively (Fig. 4A). For two of the populations that evolved in the presence of TOB+BH, a significant decrease in basal ROS production was observed (p<0.05) (Fig. 4A). These were also the populations in which mutations in BCAL2631 and/or BCAM0965 (thought to co-regulate TCA activity) were observed. When ROS levels were determined after exposure to TOB, increased ROS production was observed for one of the control populations while no significant difference was observed for any of the populations evolved in the presence of TOB alone (p<0.05) (Fig. 4B). All populations evolved in the presence of TOB+BH showed reduced ROS levels compared to the start culture after treatment with TOB; this difference was significant (p<0.05) for two lineages (Fig. 4B). Homologues of the putative ABC transporter BCAL0296, mutated in most evolved treated populations (but not in the controls), have been characterised in other bacteria. In *Bradyrhizobium* sp. the transporter homologue BclA is involved in protection against stress by antimicrobial peptides (42). BclA has multidrug transport activity and is involved in uptake of peptide-derived/peptide-like compounds, including bleomycin (43). The homologue in *Mycobacterium tuberculosis* is involved in uptake of vitamin B_12_ and bleomycin (44). It is therefore possible that BCAL0296 can import TOB into the cytoplasm under the experimental conditions used. To investigate this, we used a flow cytometry based assay to determine uptake of BODIPY-conjugated TOB. While there are no statistically significant differences when all nine groups (three lineages, three treatments) are compared (Fig. S6), differences in TOB uptake become apparent between treatments when data for the three treatments are averaged over the different lineages (Fig. 6), with TOB uptake significantly higher in the evolved control lines than in the evolved lines exposed to TOB (p = 0.045) or TOB+BH (p<0.001). No significant difference was observed between the TOB and the TOB+BH exposed cultures. The results from this assay show that there is a significantly higher fraction of the population positive for BODIPY-TOB in evolved cultures without a mutation in BCAL0296 (i.e. the controls), than in evolved cultures with a mutation in BCAL0296 (i.e. the ones exposed to TOB or TOB+BH) (Fig. 5). These data suggest that BCAL0296 is involved in TOB import in *B. cenocepacia* and that the mutations occurring after repeated exposure to TOB or TOB+BH contribute to the reduced antimicrobial activity of TOB observed.

**Fig 4.**
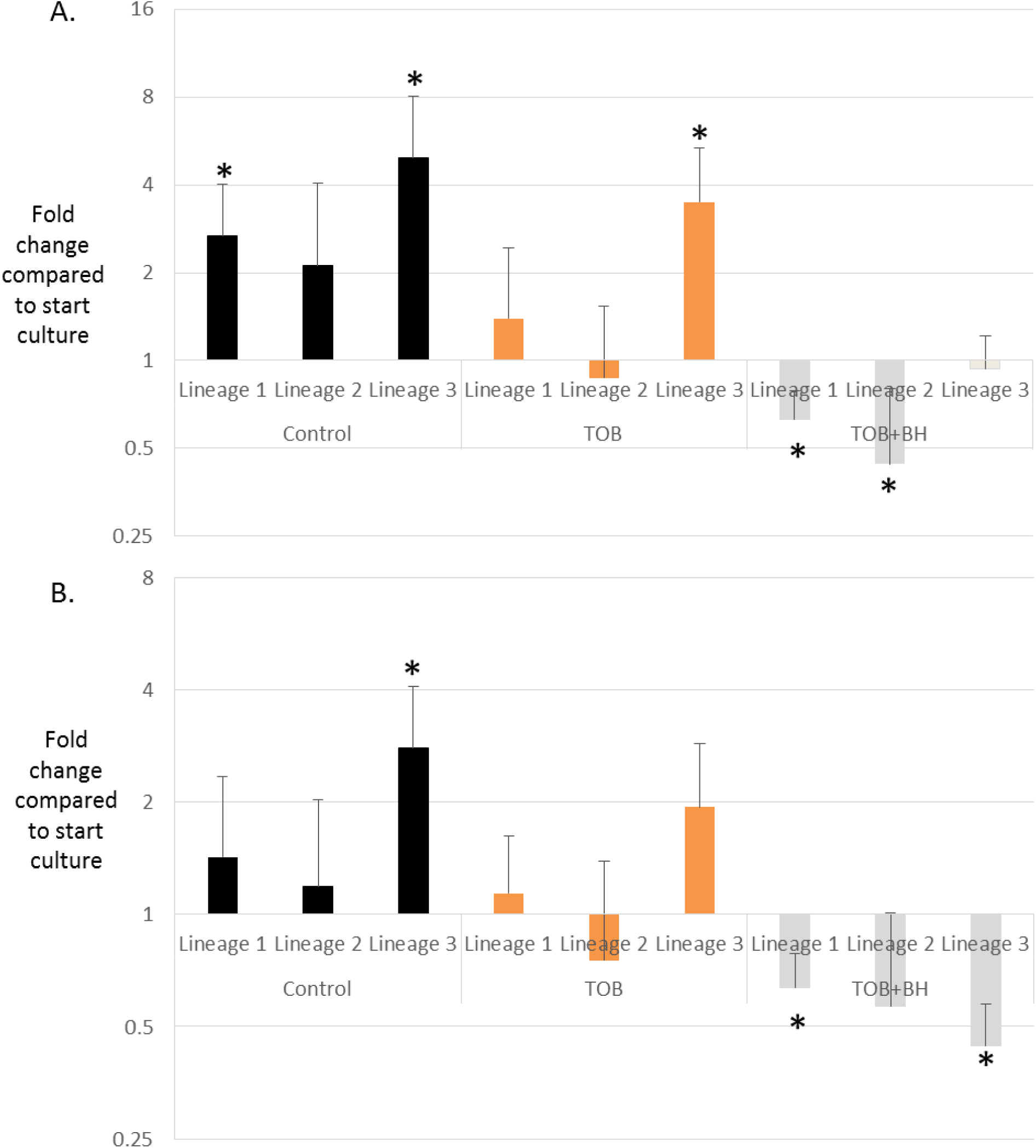
A. Basal ROS levels in evolved populations (relative compared to start culture). B. ROS levels in evolved populations after exposure to 4xMIC TOB (relative compared to start culture). Data shown are averages and error bars represent standard error (n = 3 x 5). Statistically significant differences (p < 0.05) are indicated by an asterisk.

**Fig 5.**
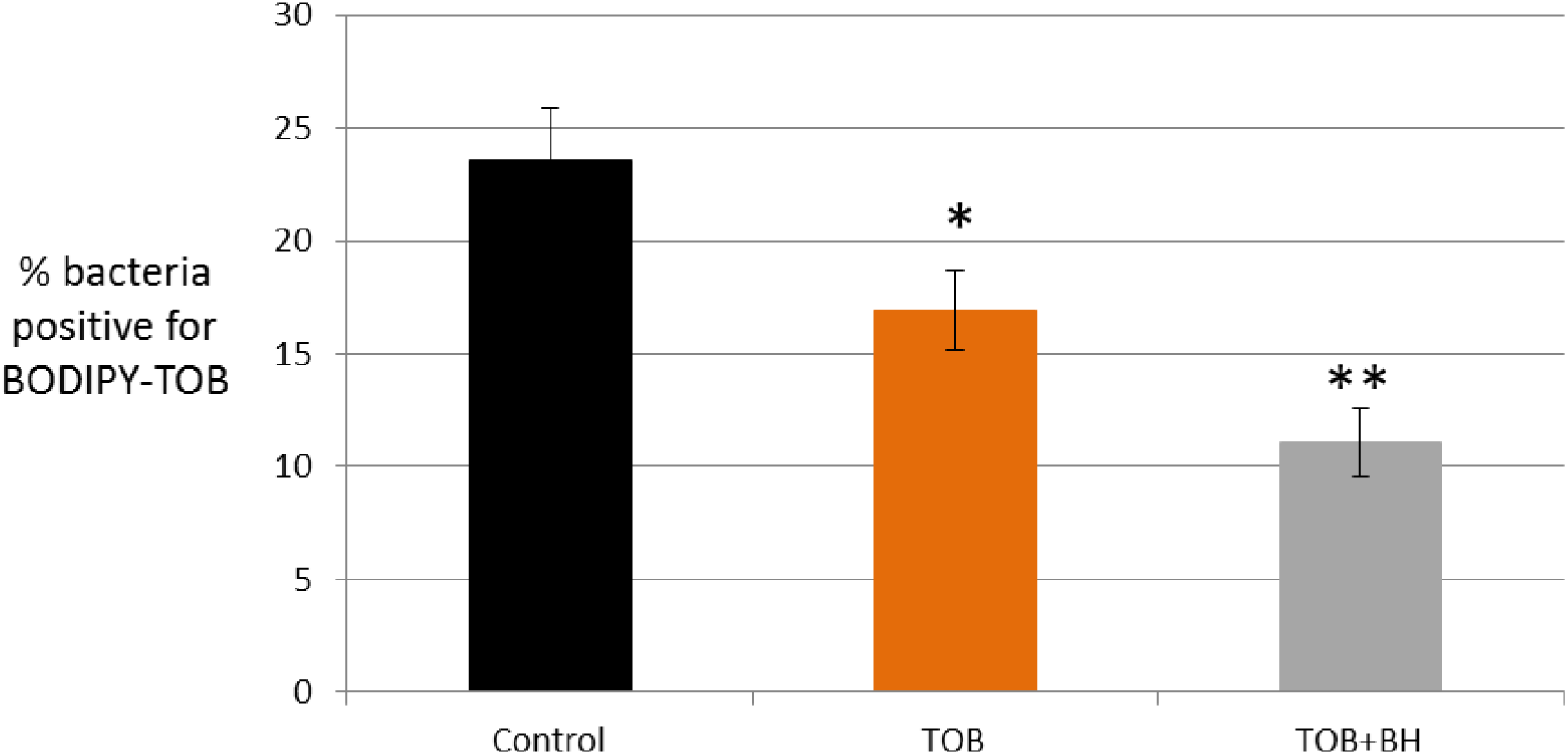
Differences in TOB uptake between treatments (based on averaged data over the different lineages). Data shown are average, error bars indicate standard error (n=24). ^*^, p < 0.05; ^**^, p < 0.001

## Conclusion

In the present study we demonstrate that during experimental evolution *in vitro*, *B. cenocepacia* J2315 biofilms gradually but quickly become less susceptible to TOB, and that evolution towards reduced susceptibility occurs significantly faster with the combined TOB+BH treatment. Many genetic changes were observed in the evolved populations exposed to the combination of TOB and BH, and some point to modifications in metabolism as a mechanism underlying the reduced susceptibility. The reduced levels of ROS (both basal levels and levels induced after exposure to TOB) observed in the lineages treated with TOB+BH point in the same direction. In addition, most lineages exposed to TOB or TOB+BH had mutations in BCAL0296, encoding an ABC transporter. Cells from populations in which BCAL0296 was mutated were more likely to accumulate lower levels of TOB intracellularly, providing an additional explanation for the reduced susceptibility of these evolved lineages. Although some genetic changes were found in multiple evolved populations, different lineages exposed to the same treatment appeared to have used different evolutionary trajectories to counteract the potentiating activity of BH. Our results indicate that resistance to potentiators can develop in multiple ways and this might limit their clinical applicability. Finally, our data demonstrate that experimental evolution combined with high-throughput sequencing can indeed identify the genetic changes behind reduced susceptibility, and allows to identify hitherto unknown genes of interest likely involved in *B. cenocepacia* biofilm resistance and tolerance (45).

## Materials and methods

### Strains and culture conditions

*B. cenocepacia* J2315 (LMG 16656) was stored at −80°C using Microbank vials (Prolab Diagnostics, Richmond Hill, ON, Canada) and subcultured at 37°C on Trypton Soy agar (TSA; Lab M, Lancashire, UK). Overnight cultures were grown aerobically in Mueller Hinton broth (MHB; Lab M) at 37°C.

### Reagents

Tobramycin (TOB; TCI Europe, Zwijndrecht, Belgium) was dissolved in physiological saline (PS) (0.9 % w/v NaCl) (Applichem, Darmstadt, Germany), filter sterilized (0.22 μm Whatman, Dassel, Germany) and stored at 4°C until use. Stock solutions of BH (Sigma-Aldrich, Bornem, Belgium) were prepared in dimethyl sulfoxide (DMSO; Sigma-Aldrich) and diluted in PS prior to use.

### Biofilm formation on beads

The set-up for biofilm formation was inspired by that reported by Traverse et al. (26). Cryobeads from Microbank vials (Prolab Diagnostics) were used as substrates for biofilm formation. The beads were rinsed with PS prior to use to remove the cryopreservative present in the Microbank vials. This was achieved by adding 1 ml PS, vortexing the vial, removing the PS and repeating this three times. Six beads were then transferred to the wells of a 24-well microtiter plate (MTP, SPL Lifescience, Korea) and one ml of a diluted overnight culture of *B. cenocepacia* J2315 (containing approximately 5 x 10^7^ colony forming units (CFU) per ml) was used as inoculum. The MTP was statically incubated at 37°C for 24 hours. To evaluate the ability of *B. cenocepacia* J2315 cells to form mature biofilms on the beads, Live/Dead staining (LIVE/DEAD BacLight bacterial viability kit, Thermo Fischer Scientific, Invitrogen, Carlsbad, CA, USA) was performed after 24 hours of biofilm formation. The biofilms on the beads were visualized using an EVOS FL Auto Cell Imaging System (Thermo Fischer Scientific, Waltham, MA, USA) (Syto9: λ_ex_ = 470/22 nm, λ_em_ = 510/42 nm; propidium iodide: λ_ex_ = 531/40 nm; λ_em_ = 593/40 nm).

### Evolution experiment

To evaluate the influence of repeated treatments on biofilm susceptibility, cells were exposed to 15 cycles of biofilm formation (24 h), treatment (24 h), and planktonic regrowth (48 h) (Fig. 1). The planktonic regrowth step was included to generate a sufficiently high number of cells to set up a new biofilm for the next cycle. Biofilms were treated with PS (untreated control), TOB alone (at a concentration of 768 μg/ml which equals 3 times the minimal inhibitory concentration [MIC]), and TOB in combination with BH (250 μM). The concentration of TOB and BH was selected based on preliminary experiments: the concentrations used in the present study lead to a significant reduction in cell numbers compared to the untreated control, but not complete eradication, so that regrowth in the following cycles can occur. Three independent experiments (designated as lineages) were set up for each condition, i.e. TOB (tobramycin), TOB+BH, and an untreated control. The three lineages were started from three different overnight start cultures. Biofilms were grown as described above and after 24 hours the beads were rinsed with PS and treated with TOB or a combination of TOB + BH. After 24 hours of treatment at 37°C, the supernatant was removed, and the beads were rinsed with PS. Each well contained 6 beads: two beads were transferred to Eppendorf tubes containing 8% dimethyl sulfoxide (DMSO; Sigma-Aldrich) in MH for storage at −80°C, while the four remaining beads were transferred to a Falcon tube containing 8 ml MH medium. Sessile cells were detached from the beads by three cycles of vortexing (1 min, Vortex-Genie 2, Scientific Industries Inc., Bohemia, NY, USA) and sonicating (1 min; Branson 3510, Branson Ultrasonics Corp, Danbury, CT, USA). Six ml of this bacterial suspension was transferred to another tube and was incubated for 48 h for regrowth, while shaking at 250 rpm at 37°C (KS 4000i control, IKA Works, Wilmington, NC, USA). The remaining 2 ml was used to determine the number of surviving cells per bead (CFU/bead) by plating.

### Determination of the minimal inhibitory concentration (MIC) and minimum bactericidal concentration (MBC)

To verify if possible changes in susceptibility over time were due to increased resistance towards TOB, the MIC and MBC for TOB was determined for the start and end population. MICs were determined according to the EUCAST broth microdilution assay using flat-bottom 96-well microtiter plates (MTP; SPL Lifescience, Korea) (46). The MIC was defined as the lowest concentration with a similar optical density as uninoculated growth medium. Absorbance was measured at 590 nm with a multilabel MTP reader (EnVision, Perkin Elmer LAS, Waltham, MA). All MIC determinations were performed in duplicate. The MBC was determined by plating the suspension used for the MIC test and the MBC was the lowest concentration that did not allow recovery of colonies following 48h incubation at 37°C.

### Determination of the number of persisters in tobramycin-exposed *B. cenocepacia* J2315 biofilms

To determine whether the evolutionary changes affected persistence, the number of persisters surviving TOB treatment was compared between biofilms formed by the start and evolved cultures. Biofilms were grown in 96 well microtiter plates as described previously (47) and exposed for 24 h to TOB in a concentration of 4 × MIC (1024 μg/ml) (39). Briefly, an inoculum suspension containing 5 × 10^7^ CFU/ml was added to the wells of a round bottomed 96 well microtiter plate. Following 4 h of adhesion, the supernatant was removed, and the plates were rinsed with PS. Subsequently, 100 μl of fresh MHB was added, and the plates were further incubated at 37 °C. After 24 h, the supernatant was removed and 120 μl of a TOB solution in PS or 120 μl PS (= control) was added. After 24 h, cells were harvested by vortexing and sonication (2 × 5 min) (Branson 3510, Branson Ultrasonics Corp, Danbury, CT) and quantified by plating on LBA. Ten wells were included per strain, and the experiment was repeated twice. (*n* = 3).

### Measurement of ROS levels

To investigate whether there were differences in the production of reactive oxygen species (ROS) between the start culture and the evolved lineages, ROS was measured in treated and untreated start and evolved cultures. To measure ROS planktonic cultures were exposed to 2’,7’-dichlorodihydrofluorescein diacetate (H2DCFDA) in a final concentration of 10 μM in LB Broth (40). After 45 min of incubation protected from light, cells were washed with PBS and treated with TOB in a concentration of 4 x MIC or pH-matched phosphate buffered saline (PBS) (= untreated control solution with the same pH as the antibiotic solution) for 24 h. Fluorescence (λ excitation = 485 nm, λ emission = 535 nm) was measured using an Envision plate reader. Autofluorescence of bacterial cells incubated without the probe and background fluorescence of the buffer solutions was measured and taken into account when calculating the net fluorescence. For the planktonic cultures an overnight culture was diluted to an optical density of 0.1 (approximately 10^8^ cells/ml). After an additional 24 h of growth in a shaking warm water bath, cell suspensions with an optical density of 1 (approximately 10^9^ cells/ml) were transferred to falcon tubes and centrifuged for 9 min at 3634 rcf. Cells were resuspended in fresh medium with or without dye to measure ROS. Five wells were included per condition and the experiment was repeated twice (n = 3 x 5).

### Genome sequencing and data analysis

After planktonic regrowth of the cells, DNA was extracted using a modified bead-beater protocol, adapted from Mahenthiralingam et al. (48). RNase-treated DNA was then quantified using the BioDrop μLITE (BioDrop, Cambridge, UK). Genomic DNA from the start culture and all evolved cultures obtained after 15 cycles were sequenced. Libraries were prepared using the NEBNext kit from Illumina, and sequenced either on an Illumina Nextseq 500 or HiSeq 4000, generating 150 bp paired-end reads (Table S3). The experimental protocols and the raw sequencing data of all samples can be found in ArrayExpress under the accession number E-MTAB-6236. Sequenced reads were quality trimmed (error probability limit 0.05) and mapped to the *B. cenocepacia* J2315 reference genome (34) using CLC Genomics Workbench version 11.0.1. (Qiagen, Aarhus, Denmark) with a cut-off of 80% for similarity and 50% mapped read length. Mapping parameters were: match score 1, mismatch cost 2, insertion and deletion cost 3. More than 98.6% of reads mapped to the reference genome for all samples (Table S3). The un-mapped reads were *de-novo* assembled in CLC Genomics Workbench, but no contigs with a coverage >10% of the average coverage of the respective sample were found and gene acquisition was therefore excluded. In CLC Genomics Workbench, the InDels and Structural Variants tool was used to detect insertions and deletions, with a p-value threshold of 0.0001. The output was manually screened on mapping patterns of un-aligned read ends and only entries with a single breakpoint and identical sequences in the un-aligned read ends were reported. The consensus sequence of the un-aligned read ends was then used to confirm the deletion or to identify the nature of the inserted sequence. A larger insertion sequence cannot be fully deduced in this manner, but only a certain type of transposase which was already present multiple times in the *B. cenocepacia* J2315 reference genome was detected: *Burkholderia cepacia* insertion element IS*407*. Both consensus sequences at insertion breakpoints were consistent with either end of this transposase, it was therefore concluded that the insertion consisted of only that transposase. The Basic Variant Detection tool was used to detect Single Nucleotide Polymorphisms (SNPs) with a minimum coverage of 10 and a reference-to-variant ratio of 35%. This was the lowest cut-off that allowed to clearly distinguish true SNPs from sequencing errors. All SNPs were then manually screened for false positives in regions containing repetitive sequences or hairpins, which caused poor mapping. The function of the genes that acquired mutational changes was determined using the Conserved Domain database (49) and *Burkholderia* Genome Database (50).

### qPCR

Cultures from cycle 15, control lineage 3 and Tob lineage 2, were cultivated in MHB in a shaking incubator at 150 rpm for 6 to 10 hours. Late log phase cultures were harvested at a density of 1 - 1.3 x 10^9^ CFU/ml and stationary phase cultures at a density of 3 - 4.5 x 10^9^ CFU/ml. Cell pellets were frozen at −80°C and RNA was extracted within one week of harvest, using the RiboPure bacteria kit (Thermo Fisher, Rochester, NY, USA), according to the standard protocol, including DNase treatment. RNA was quantified with the BioDrop μLITE. cDNA was generated with the High Capacity cDNA Reverse Transcription kit (Applied Biosystems, Foster City, CA, USA), from 500 ng of RNA. qPCR was performed in a CFX96 Real-Time System C1000 Thermal Cycler (Bio-Rad, Hercules, CA, USA) using GoTaq qPCR Master Mix (Promega, Madison, WI, USA). Cq values were normalised against a previously-validated control gene (*rpo*D, BCAM0918) (51, 52). Fold changes were calculated compared to a standard (mix of all cDNAs in experiment) and log-transformed. Primers are listed in Table S4.

### Growth curves

Growth curves were determined in MHB. 200 μl/well of a 5 x 10^5^ CFU/ml inoculum was added in triplicates to a round-bottom MTP and the absorbance at 590 nm was measured in a microplate reader (Envision, Perkin Elmer, Shelton, CT, USA), every 30 minutes for 50 hours. The experiment was repeated 3 times and representative curves are shown.

### Determination of CepI activity in the presence of BH

*B. cenocepacia* CepI was expressed in *Escherichia coli* BL21(DE3) cells and purified according to the procedure previously described (53). Enzymatic activity was determined by a spectrophotometric assay, according to Christensen *et al*. (54), which measures the *holo*-ACP formation by titrating the release of the free thiol of ACP with dichlorophenylindophenol (DCPIP; ε= 19100 M^-1^ cm^-1^). Measurements were performed at 37 °C, in a final volume of 100 μl, using an Eppendorf Biospectrometer. The standard reacion mixture contained 50 mM Hepes pH 7.5, 0.005% Nonidet P-40, 0.13 mM DCPIP, 70 μM Octanoyl-ACP (C8-ACP) (55, 56) and 4 μM CepI; the reactions were started by addition of 40 μM S-adenosyl methionine (SAM), after pre-incubation for 10 min. CepI inhibition was initially screened at 200 μM of BH (dissolved in DMSO). CepI inhibition was initially screened at 200 μM (dissolved in DMSO) and subsequently the IC_50_ was determined by measuring enzyme activities in presence of different BH concentrations, and fitting data according to equation (1):

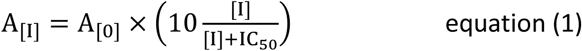

where A_[I]_ is the enzyme activity at BH concentration [I] and A_[0]_ is the enzyme activity without BH. All measurements were performed in triplicate.

### Quantification of intracellular tobramycin levels

BODIPY-labelled tobramycin was synthesized as previously described (57). Bacteria were grown under biofilm-forming conditions for 4 h in MHB in the presence of 0.75 μg/mL BODIPY-tobramycin at 37 °C. Following 4 h of biofilm formation, the biofilm was rinsed to remove extracellular tobramycin, homogenized and subjected to flow cytometry analysis (Attune NxT, Life Technologies). The bacterial population was delineated based on the forward and side scatter signal, and a threshold was set to exclude non-cellular particles and cell debris. BODIPY-tobramycin that associated with bacterial cells was determined through excitation with a 488 nm laser. Fluorescence emission was detected through a 530/30 bandpass filter. Controls included bacterial biofilm cells that were not exposed to BODIPY-tobramycin (negative control) or to incremental levels of tobramycin to determine the concentration at which saturation was obtained. Based on the negative control and the concentration of tobramycin where maximal population saturation was obtained, negative and positive flow cytometry gates were determined respectively. At least 10,000 bacteria were analysed per sample.

### Statistical analysis

To determine whether the observed variations in survival over time for the different treatments were statistically significant, a linear mixed-effect model (LMEM) was used. The model uses log(CFU/bead) as the dependent variable and cycle, treatment, lineage and their two- and three-way interactions as fixed effects and was fit using SAS version 9.4 (SAS institute, Cary, NC, USA). To account for possible correlations between the measurements over cycles, a compound symmetry variance covariance structure was used. All interaction effects that were not significant were excluded from the model. When an interaction was significant, this was considered as the fixed effect to evaluate differences in treatment effect. Per lineage, treatments were compared pairwise to TOB treatment using the Tukey adjustment method. Assumptions associated with the LMEM were checked based on residuals from the fitted final model (Table S2).

Other data sets were analysed using SPSS version 25 software (SPSS, Chicago, IL, USA). The Shapiro-Wilk test was used to verify the normal distribution assumption of the data. Normally distributed data were analysed using a one-way ANOVA, while non-normally distributed data were analysed with a Kruskal-Wallis 1-way ANOVA. P-values smaller than 0.05 were considered statistically significant.

